# Diffuse TBI-induced expression of anxiety-like behavior coincides with altered glutamatergic function, TrkB protein levels and region-dependent pathophysiology in amygdala circuitry

**DOI:** 10.1101/640078

**Authors:** Joshua A. Beitchman, Daniel R. Griffiths, Yerin Hur, Sarah B. Ogle, Caitlin E. Hair, Helena W. Morrison, Jonathan Lifshitz, P. David Adelson, Theresa Currier Thomas

**Affiliations:** Barrow Neurological Institute at Phoenix Children’s Hospital, Phoenix, AZ, USA; Dept of Child Health, University of Arizona College of Medicine-Phoenix, Phoenix, AZ, USA; Midwestern University, Glendale, AZ, USA; University of Arizona, Tucson, AZ, USA; Phoenix VA Healthcare System, Phoenix, AZ, USA; Banner University Medical Center, Phoenix, AZ, USA

## Abstract

Up to 50% of traumatic brain injury (TBI) survivors demonstrate persisting affective symptoms indicative of limbic system dysregulation, yet the pathophysiology underlying the symptoms is unclear. We hypothesize that TBI-induced pathophysiologic changes within distinct amygdala nuclei contribute to the expression of late-onset anxiety-like behavior. Adult, male Sprague-Dawley rats underwent midline fluid percussion injury or sham surgery. Anxiety-like behavior was assessed at 7 and 28 days post-injury (DPI) followed by assessment of real-time glutamate neurotransmission in the basolateral amygdala (BLA) and central nucleus of the amygdala (CeA) using glutamate-selective microelectrode arrays. In separate animal cohorts, the presence of neuropathology, astrocytosis, and microglial activation were assessed at 1, 7, and 28DPI. Protein levels of glutamatergic transporters (Glt-1, GLAST) and presynaptic modulators of glutamate release (mGluR2, TrkB, BDNF, and glucocorticoid receptors) were quantified using automated capillary western techniques at 28DPI. The expression of anxiety-like behavior at 28DPI coincided with decreased glutamate release and slower glutamate clearance in the CeA, not BLA. Changes in glutamate neurotransmission were independent of protein levels of glutamate transporters and mGluR2 receptors, neuropathology, and astrocytosis. At 1DPI, microglia in the CeA demonstrated a neuroinflammatory response. BDNF and TrkB were decreased at 28DPI in the amygdala. These data indicate that diffuse axonal injury instigates sequences of molecular, structural and functional changes in the amygdala that contribute to circuit dysregulation relevant to the expression of affective disorders. Translationally, diffuse axonal injury can influence severity and incidence of affective symptoms and should be addressed in the history of patients with affective disorders.

## Introduction

Affective disorders, including generalized anxiety disorder and post-traumatic stress disorder (PTSD), are reported in 18.1% of United States adults. Additionally, affective disorders develop and persist in up to 50% of traumatic brain injury (TBI) survivors, however, few common biological mechanisms between TBI and non-TBI patients have been elucidated ^1, 2^. The incidence of TBI continues to rise, with at least 2.5 million Americans reporting a TBI each year, costing the American health care system 76.5 billion dollars ^3^. Three-quarters of all TBIs are diffuse TBIs (dTBI) in which the signature pathology is multi-focal diffuse axonal injury with no overt pathology detected by CT or MRI ^4-6^. dTBI subsequently induces secondary sequelae that occur seconds to months following the initial injury, leading to the development of affective, cognitive, and somatic symptoms ^7, 8^. Despite clinical prevalence, the pathophysiology contributing to affective symptoms following dTBI is poorly understood, resulting in mis-diagnosis and ineffective treatments ^9-11^. Ultimately, affective symptoms impact a patient’s ability to return to work, daily function, and social interactions, and thus severely impair the quality of life for both the patient and caregivers ^12, 13^.

Clinical and preclinical data demonstrate how the amygdala changes following dTBI. In veterans, early amygdala fMRI reactivity post-injury is predictive of the development of PTSD and could contribute to the bilateral reduction of amygdala size observed in patients diagnosed with both TBI and PTSD ^14, 15^. A study in collegiate football players report a positive correlation between amygdala shape and mood states ^16^. TBI survivors with major depressive disorder had smaller lateral and dorsal prefrontal cortex, which mediates ventral-limbic and paralimbic pathways and influences amygdala circuitry ^17^. Diffuse and focal experimental models of TBI also identify alterations in amygdala circuitry, including increased neuronal hyperexcitability and GABA production proteins in the absence of overt neuropathology ^18, 19^. In addition, pyramidal and stellate neurons in the basolateral amygdala (BLA) demonstrate increased complexity distal and proximal to the soma as early as 1 day post-injury (DPI) and lasted until at least 28DPI indicating BLA-central nucleus of the amygdala (CeA) circuit reorganization after dTBI ^20^.

Affective symptoms are controlled by limbic system circuitry with evidence that the direct stimulation of glutamatergic neurons originating in the BLA and projecting into the CeA produces reversible anxiolytic symptoms ^21 22^. BLA neurons are 90% glutamatergic, highlighting the role of glutamate neurotransmission in displays of affective behavior ^23, 24^. Glutamate neurotransmission is regulated by astrocytes and microglia processes surrounding the synapse ^25^. Glutamate transporters, Glt-1 and GLAST (located on astrocytes), rapidly remove glutamate from the extracellular space, restricting glutamate to the synaptic cleft. ^26, 27^. When spillover does occur, feedback through the metabotropic glutamate receptor 2 (mGluR2) can reduce presynaptic release of glutamate ^28^. Decreased levels of brain derived neurotropic factor (BDNF), tropomyosin-related kinase B (TrkB) receptors, and glucocorticoid receptors (GluR)-commonly reported with affective disorders - have also been reported to attenuate presynaptic glutamate release ^29-31^. In addition, microglia also express glutamate receptors and can respond to and mediate glutamate signaling ^25^.

Clinical and experimental studies report increased prevalence of anxiety symptoms, stress resilience, and mood disorders over time after focal and diffuse TBI, with complementary experiments indicating a regulatory role of the amygdala in the genesis of these abnormal affective symptoms. Yet, the pathophysiological processes, including circuit function, structure, and molecular modulators have not been elucidated. We hypothesize that pathophysiologic changes within distinct amygdala nuclei precede or coincide with TBI-induced anxiety-like behavior. In a well-established model of diffuse axonal injury (DAI) in rodents, we assessed the expression of anxiety-like behavior, immediately followed by *in vivo* amperometric recordings of glutamate neurotransmission in the BLA and CeA. Pathological and molecular analyses of nuclei-specific changes were completed for mechanistic evaluation.

## Materials and Methods

### Animals

A total of 69 adult, male Sprague-Dawley rats (weights 279-420 grams; age 3-4 months) (Envigo, Indianapolis, IN) were used in these experiments. (29 rats for amperometry, 22 for histology, and 18 rats for protein assays). Upon arrival, rats were given a 1 week acclimation period, housed in normal 12-hour light/dark cycle (Red light: 18:00 to 06:00) and allowed access to food and water *ad libitum* (Teklad 2918, Envigo, Indianapolis, IN). Rats were pair housed according to injury status (i.e. injured housed with injured) throughout the duration of the study. All procedures and animal care were conducted in compliance with an approved Institutional Animal Care and Use Committee protocol (13-460) at the University of Arizona College of Medicine-Phoenix which is consistent with the National Institutes of Health (NIH) Guidelines for the Care and Use of Laboratory Animals.

Rats undergoing neurological assessment were exposed to human contact and handled for a total of fifty minutes over a period of 7 days. Rats aged out to 28DPI received an additional thirty minutes of handling over a period of 3 days immediately prior to testing to reacclimatize to human contact. At 7or 8DPI or 28 or 29DPI, injured and sham rats underwent neurological assessment followed by *in vivo* amperometric recordings. For the remainder of the manuscript, DPI for the amperometric and behavioral outcome measures at 7/8 and 28/29DPI will be abbreviated as 7DPI and 28DPI. For behavioral analysis and subsequent amperometric recordings, a total of 17 rats were aged to the one-week time point (11 injured; 6 sham) and 17 rats were aged to one-month (10 injured; 7 sham) (experimental design; Figure 1A).

**Figure 1:**
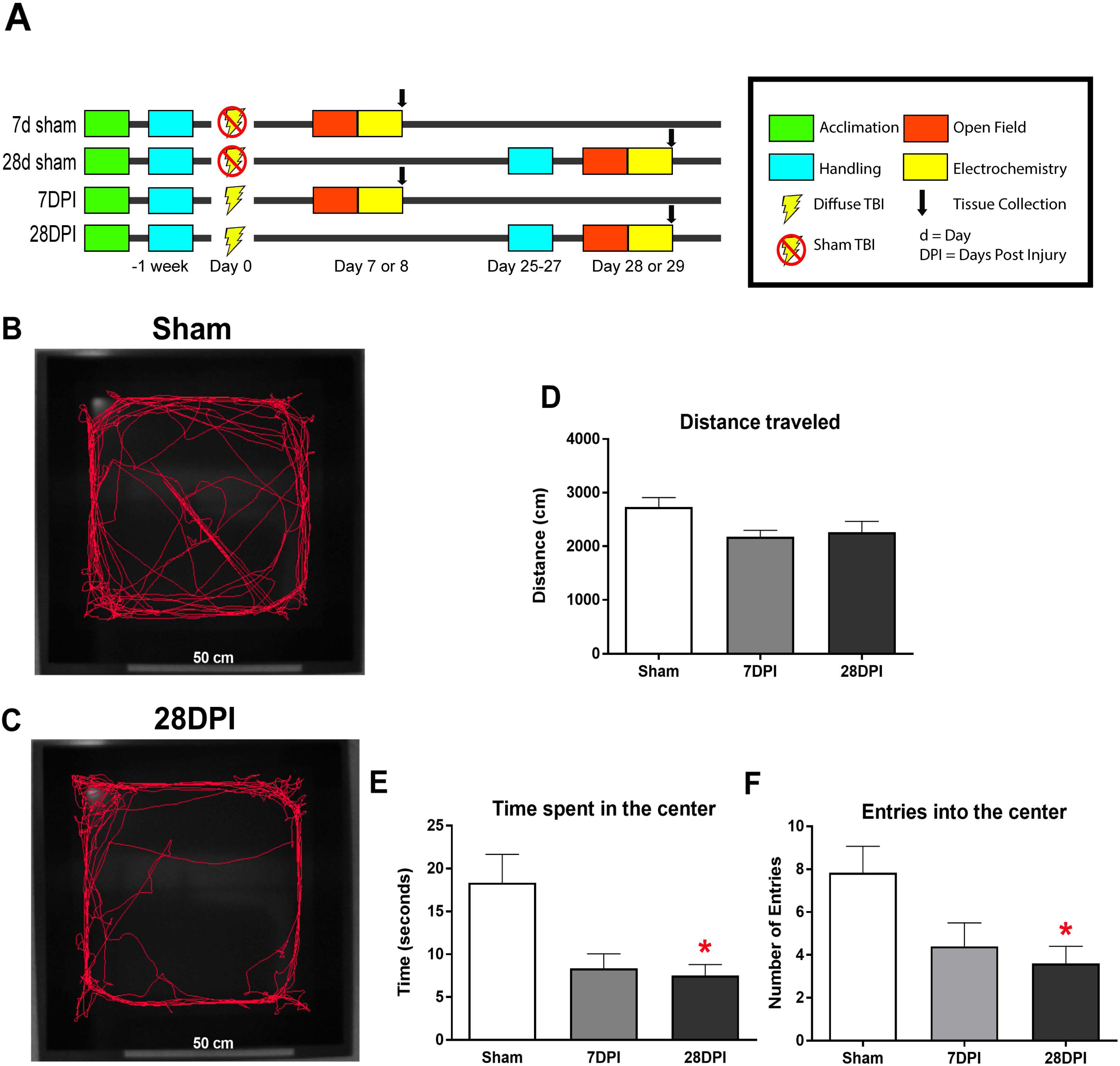
dTBI induces the expression of anxiety-like behavior. **A)** Study design for behavioral and amperometry studies. **B)** Representative video tracking data from Ethovision^®^ software during the first 5 minutes of open field testing from sham. **C)** Representative video tracking data from Ethovision^®^ software during the first 5 minutes of open field testing from 28DPI rats. **D)** Distance traveled is similar between 7DPI, 28DPI, and sham [One-Way ANOVA F(2,29)=3.23; p=0.05]. **E)** 28DPI rats spent significantly less time in the center of the open field [Kruskal-Wallis H=6.711; p<0.05; Dunn’s post-hoc comparison p<0.05; Mann-Whitney U post-hoc p<0.05, r=0.48]. **F)** 28DPI rats made significantly fewer entries into the center of the open field when compared to sham [Kruskal-Wallis H=6.83; p<0.05; Dunn’s post-hoc comparison p<0.05; Mann-Whitney U post hoc p<0.05, r=0.46]. Bar graphs represents mean ± SEM. *n*=10-12 for each group.

### Midline Fluid Percussion Injury (mFPI)

#### Surgical procedure

Midline FPI surgery was carried out similarly to previously published methods from this laboratory ^32^. Rats were randomized into either injured or sham groups following acclimation to the vivarium facility and exposure to human contact. Briefly, rats were anesthetized with 5% isoflurane in 21% O_2_ and placed into a stereotaxic frame (Kopf Instruments, Tujunga, CA) with a nose-cone that maintained 2.5% isoflurane for the duration of the procedure. A 4.8 mm circular craniotomy was performed centered on the sagittal suture midway between bregma and lambda carefully ensuring that the underlying dura and superior sagittal sinus were not punctured or disturbed. An injury hub created from the female portion of a 20-gauge Luer-Loc needle hub was cut, beveled and placed directly above and in-line with the craniectomy site. A stainless-steel anchoring screw was then placed into a 1 mm hand-drilled hole into the right frontal bone. The injury hub was affixed over the craniectomy using cyanoacrylate gel and methyl-methacrylate (Hygenic Corp., Akron, OH) and filled with 0.9% sterile saline. The incision was then partially sutured closed on the anterior and posterior edges with 4.0 Ethilon sutures and topical lidocaine and antibiotic ointment were applied. Rats were returned to a warmed holding cage and monitored until ambulatory (approximately 60-90 min).

#### Injury induction

Approximately two-four hours following surgical procedures and the return of ambulation, rats were re-anesthetized. The hub was filled with 0.9% sterile saline and attached to the male end of a fluid percussion device (Custom Design and Fabrication, Virginia Commonwealth University, Richmond, VA). After return of the pedal withdrawal reflex, an injury averaging 2.19 atm was administered by releasing the pendulum (16.5 degrees) onto the fluid-filled cylinder. Shams were attached to the fluid percussion device, but the pendulum was not released. Immediately after the administration of injury, the injury hub was removed *en bloc* and rats monitored for the presence of apnea, fencing response, and the return of righting reflex. Then, rats were briefly re-anaesthetized to inspect the injury site for hematoma, herniation, and dural integrity. The injury site was then stapled closed and topical lidocaine and antibiotic ointment were applied. Injured rats had a righting reflex time ranging from 5 to 12 min (average 7:43 ± 0:07 minutes) with 100% of brain-injured rats demonstrating the fencing response. Rats were placed in a clean, warmed holding cage, and monitored for at least one hour following injury before being returned to the vivarium where post-operative evaluations continued for three to five DPI. Staples for rats with a 28-day time point were removed at seven DPI.

### Neurological assessments

#### Open field

Each rat was placed into the center of a novel black open field (measuring 69 cm by 69 cm) facing the same direction at least one hour following the light cycle change from red to white light. White overhead lights ensured equal illumination throughout the container for a total of 15 minutes. Ambient white noise (60-70 decibels) in the room mitigated any variation in noise levels. Tracking was accomplished by Ethovision ^®^ Software which mapped the rat directly overhead. Two regions were defined using Ethovision^®^, separating the open field into an outer region and an inner region measuring approximately 35 cm by 35 cm. Time spent in each zone was calculated based on the center-point of the rat. Primary outcome measures analyzed total distance traveled, time spent in the center, and entries to the center of the open field for the first 5 minutes of the task to be more representative of anxiety-like behavior ^33^.

### Microelectrode arrays

#### Enzyme-based microelectrode arrays

Ceramic-based microelectrode arrays (MEA) encompassing four platinum (Pt) recording surfaces (15 µm x 333 µm) aligned in a dual, paired configuration were prepared to measure glutamate for *in vivo* anesthetized recordings (S2 configuration; Quanteon, Nicholasville, KY). MEAs were fabricated, selected for recordings, and made glutamate sensitive as previously described ^32^. Briefly, Pt sites 1 and 2 were coated with a solution containing glutamate oxidase (GluOx), bovine serum albumin (BSA), and glutaraldehyde, enabling these sites to selectively detect glutamate levels with low limits of detection ^34^. Pt sites 3 and 4 were coated with only BSA and glutaraldehyde and served as sentinels, recording everything channels 1 and 2 recorded except for glutamate ^35^. Prior to calibration and *in vivo* recordings, all four Pt recording sites were electroplated with a size exclusion layer of 1,3-phenylendediamine (mPD) (Acros Organics, Morris Plains, NJ). GluOx converts glutamate into α-ketoglutarate and peroxide (H_2_O_2_). The H_2_O_2_ functions as a reporter molecule, traversing the mPD layer and is readily oxidized and recorded as current using the FAST-16 mkIII system (Fast Analytical Sensor Technology Mark III, Quanteon, L.L.C., Nicholasville, KY) (Supplementary Figure 1).

#### Microelectrode array calibration

On the morning of *in vivo* recordings, each MEA was calibrated *in vitro* to determine recording parameters: slope (sensitivity to glutamate), limit of detection (LOD; lowest amount of glutamate to be reliably recorded), and selectivity (ratio of glutamate to ascorbic acid). For calibration, aliquots from stock solutions were added to 40 mL of 0.05 M phosphate buffered saline (PBS) (pH 7.1-7.4; stirring; 37°C) in the following sequence: 500 µL of 20 mM ascorbic acid, three additions of 40 µL of 20 mM l-glutamate, and 40 µL of 8.8 mM H_2_O_2_ to produce a final concentration of 250 μM AA, 20, 40, and 60 μM glutamate, and 8.8 μM H_2_O_2_. A representative MEA calibration is shown in Figure 2A. For the study, a total of 48 MEAs were used with a total of 91 recording sites. Additionally, there was an overall average slope of 6.9 pA/µM, a LOD of 2.4 µM, and a selectivity of 44 to 1.

**Figure 2:**
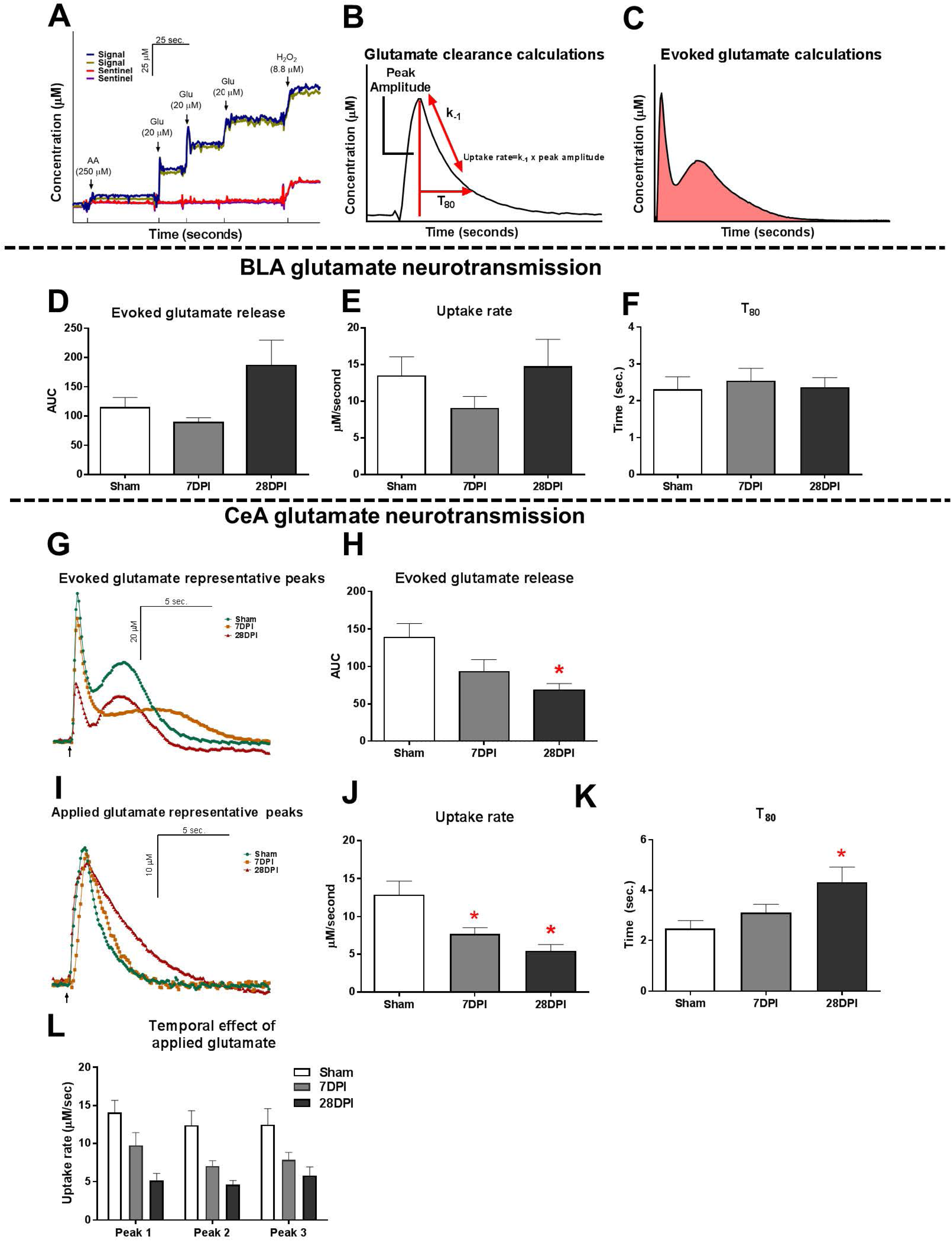
TBI induces altered glutamate neurotransmission in the CeA. **A)** Representative calibration of glutamate selective MEA. Arrows represent aliquots of solution of either 250 µM ascorbic acid (AA), 20 µM glutamate (Glu) or 8.8 µM H_2_O_2_. **B)** Calculations for total evoked glutamate release following local applications of 120 mM potassium chloride solution (KCl). **C.** Calculations for extracellular clearance of glutamate following local applications of 100 µM glutamate. **D)** No significant differences were observed in evoked glutamate release in the BLA [Kruskal Wallis H=4.63; p=0.10; *n*=8-11]. **E)** No changes in glutamate clearance were observed when assessing the uptake rate [One-way ANOVA F(2,21)=1.24; p=0.31] or **F)** T_80_ [One-way ANOVA F(2,21)=0.14; p=0.87; *n*=6-10]. **G)** Representative traces of KCl-evoked glutamate release. Baseline levels have been adjusted to show comparison amongst the three time points. Arrow represents 120 mM KCl administration. **H)** 28DPI rats had 50% less total evoked glutamate when compared to shams [One way ANOVA F(2,19)=4.74 p<0.05 Dunnett’s post-hoc p<0.05, η^2^=0.33; *n*=6-9] in the CeA. **I)** Representative traces of extracellular glutamate clearance. Baselines levels have been adjusted to show comparison amongst the three time points. Arrow represents 100 µM glutamate administration. **J)** 7 and 28DPI rats had 40% and 58% slower uptake rate, respectively [One-way ANOVA F(2,22)=7.88; p<0.01, @ 7DPI Dunnett’s post-hoc p<0.05, @ 28DPI Dunnett’s post-hoc p<0.01, η^2^=0.41; *n*=7-10]. **K)** 28DPI rats had a 43% increase in T_80_ [One-way ANOVA F(2,22)=5.00; p<0.05, Dunnett’s post-hoc p<0.01, η^2^ = 0.33; *n*=7-10]. **L)** Uptake rate remained consistent between subsequent additions of glutamate to the CeA [RM Two-way ANOVA F(2,21)=8.16; p<0.01; *n*=7-10]. Bar graphs represents mean ± SEM.

#### Microelectrode array/micropipette assembly

Following calibration, a single micropipette was attached to the MEA using the following steps to allow for the local application of solutions during *in vivo* experiments. A single-barreled glass capillary with filament (1.0 x 0.58 mm^2^, 6” A-M Systems, Inc., Sequim, WA) was pulled using a Kopf Pipette Puller (David Kopf Instruments, Tujunga, CA). Using a microscope with a calibrated reticle, the pulled micropipette was bumped against a glass rod to have an inner diameter of 7-13 µm (10.5 µm ± 0.2). Clay was used to place the tip of the micropipette between the 4 Pt recording sites. This alignment was secured using Sticky Wax (Kerr Manufacturing Co). The final measurements were the distance between the micropipette tip and the MEA surface (72 µm ± 3) and the distance between the micropipette tip and the MEA tip (498 µm ± 4).

#### Surgery for amperometric recordings

Immediately after neurological assessments, sham and brain-injured rats were anesthetized with three or four intraperitoneal injections of 25% urethane in 15-minute intervals (1.5 g/Kg; Sigma Aldrich, St. Louis, MO). Following cessation of a pedal withdrawal reflex, each rat was then placed into a stereotaxic frame (David Kopf Instruments) with nonterminal ear bars. Body temperature was maintained at 37°C with isothermal heating pads (Braintree Scientific, Braintree, MA). A midline incision was made and the skin, fascia, and temporal muscles were reflected to expose the skull. A bilateral craniectomy exposed the stereotaxic coordinates for the BLA and CeA. Dura was then removed prior to the implantation of the MEA. Brain tissue was kept moist through the application of saline soaked cotton balls and gauze. Finally, using blunt dissection, an Ag/AgCl coated reference electrode wire was placed in a subcutaneous pocket on the dorsal side of the subject ^36, 37^.

#### *In vivo* amperometric recording

Amperometric recordings performed here were done similar to previous published methods ^32, 38^. Briefly, a constant voltage was applied to the MEA using the FAST-16 mkIII recording system. *In vivo* recordings were performed at an applied potential of +0.7V compared to the Ag/AgCl reference electrode. All data were recorded at a frequency of 10 Hz, amplified by the headstage (2 pA/mV) without signal processing or filtering of the data.

Immediately prior to implantation of the MEA, the pipette was then filled with 120 mM KCl (120 mM KCl, 29mM NaCl, 2.5mM CaCl_2_ in ddH_2_O, pH 7.2 to 7.5) or 100 µM l-glutamate (100 µM l-glutamate in 0.9% sterile saline pH 7.2-7.6). Concentrations for both solutions have been previously shown to elicit reproducible KCl-evoked glutamate release or exogenous glutamate peaks ^32, 39^. Solutions were filtered through a 0.20 µm sterile syringe filter (Sarstedt AG & Co. Numbrecht, Germany) and loaded into the affixed micropipette using a 4-inch, 30-gauge stainless steel needle with a beveled tip (Popper and Son, Inc, NY). The open end of the micropipette was connected to a Picospritzer III (Parker-Hannin Corp., General Valve Corporation, Mayfield Heights, OH) with settings to dispense nanoliter quantities of fluid over a 1 second period using the necessary pressure of nitrogen (inert) gas using a dissecting microscope (Meiji Techno, San Jose, CA) with a calibrated reticle in the eyepiece ^40, 41^.

The MEA-micropipette was localized in the BLA (A/P: ±2.4 mm; M/L: ±5.1 mm; D/V: −8.0 mm) and CeA (A/P: ±2.4 mm; M/L: ±3.9 mm; D/V: −8.0 mm) using coordinates from George Paxinos and Charles Watson Rat Brain Atlas (6^th^ Edition) through use of the Dual Precise Small Animal Stereotaxic Frame (Kopf, model 962). Glutamate and KCl-evoked measures were recorded in both hemispheres in a randomized and balanced experimental design to mitigate possible hemispheric variations or effect of anesthesia duration.

#### KCl-evoked release of glutamate analysis parameters

Once the electrochemical signal had reached baseline, 120 mM KCl was locally applied (BLA: 110 nl ± 8; CeA: 105 nl ± 4) to produce an evoked glutamate release. Additional ejections of KCl were completed at 2-minute intervals and were volume matched at the time of administration. Criteria for analysis required that the peak with the largest amplitude was acquired from the first local application of KCl. This ensured that the data chosen were most representative of the maximum glutamate released within the surrounding neuronal tissue. Primary outcome measures were area under the curve as a proxy to investigate the total glutamate release capable of the recorded region. For a diagrammatic representation of these calculations, see Figure 2B.

#### Glutamate clearance analysis parameters

Once the baseline was reached and maintained for at least 2 minutes (10-20 minutes), 100 µM glutamate was locally applied into the extracellular space (BLA: 73 nl ± 17; CeA: 78 nl ± 15). Exogenous glutamate was released at 30 second intervals and amplitude was matched at the time of administration. In analysis, three peaks were selected based on a predetermined amplitude range of 15 to 25 µM to ensure that data chosen were most representative of the glutamate clearance of similar volume and amplitude in accordance with Michaelis-Menton kinetics clearance parameters. The parameters for the 3 peaks were then averaged to create a single representative value per recorded region per rat. Primary outcome measures analyzed the uptake rate and the time taken for 80% of the maximum amplitude of glutamate to clear the extracellular space (T_80_). The uptake rate was calculated using the uptake rate constant (k_-1_) multiplied by the peak’s maximum amplitude, thus controlling for any variation between the amount of applied glutamate between peaks. For a diagrammatic representation of these calculations, see Figure 2C.

#### MEA placement verification

Immediately following *in vivo* anesthetized recordings, rats were transcardially perfused with PBS followed directly by 4% paraformaldehyde (PFA). Brains were cryoprotected and sectioned at 40 µm sections to confirm MEA electrode placement. Of these, 1.96% of electrode tracts were excluded due to inaccurate placement. Representative image shown in Supplementary Figure 2.

### Microglia skeleton analysis

#### IBA1 staining

At 1, 7, and 28DPI, sham and brain-injured rats (n=3/time point) were administered a lethal dose of Euthasol^®^ (a sodium pentobarbital mixture, 200 mg/kg, i.p., Virbac AH, Inc.) and were transcardially perfused with PBS, followed by a fixative solution containing 4% paraformaldehyde. Brains were then shipped to Neuroscience Associates Inc. (Knoxville, TN) where they were embedded into a single gelatin block (MultiBrain^®^ Technology, NeuroScience Associates, Knoxville, TN). Forty micron thick sections were taken in the coronal plane and wet-mounted on 2%-gelatin-subbed slides before being stained with ionized calcium binding adaptor molecule (Iba1) primary antibody and 3,3′-Diaminobenzidine (DAB) visualization (NeuroScience Associates, Knoxville, TN) to identify all microglia.

Photomicrographs of the BLA and CeA were taken using a Zeiss microscope (Imager A2; Carl Zeiss, Jena, Germany) in bright-field mode with a digital camera using a 40x objective. One digital photomicrograph in each region was acquired from the left and right hemisphere respectively for each time point across three coronal sections for each mFPI and sham rat, for a total of 6 images per rat. A computer-aided skeletal analysis method was used to quantify morphological remodeling of ramified microglia after experimental diffuse brain injury ^42^.

#### Skeleton analysis

Microglia were analyzed by an investigator blinded to injury status using computer-aided skeleton analysis as previously published ^42^. Briefly, photomicrographs were converted to binary images which were skeletonized using ImageJ software (National Institutes of Health, https://imagej.nih.gov/ij/)(Supplementary Figure 3). The Analyze Skeleton Plugin (developed by and maintained here: http://imagej.net/AnalyzeSkeleton31) was applied to the skeletonized images, which tags branches and endpoints and provides the total length of branches and total number of endpoints for each photomicrograph. Cell somas were manually counted by 2 investigators and averaged for each photomicrograph. The total branch length and number of process endpoints were normalized to number of microglia cell somas per image. Data from the 6 images were averaged to a single representative measure per animal.

**Figure 3:**
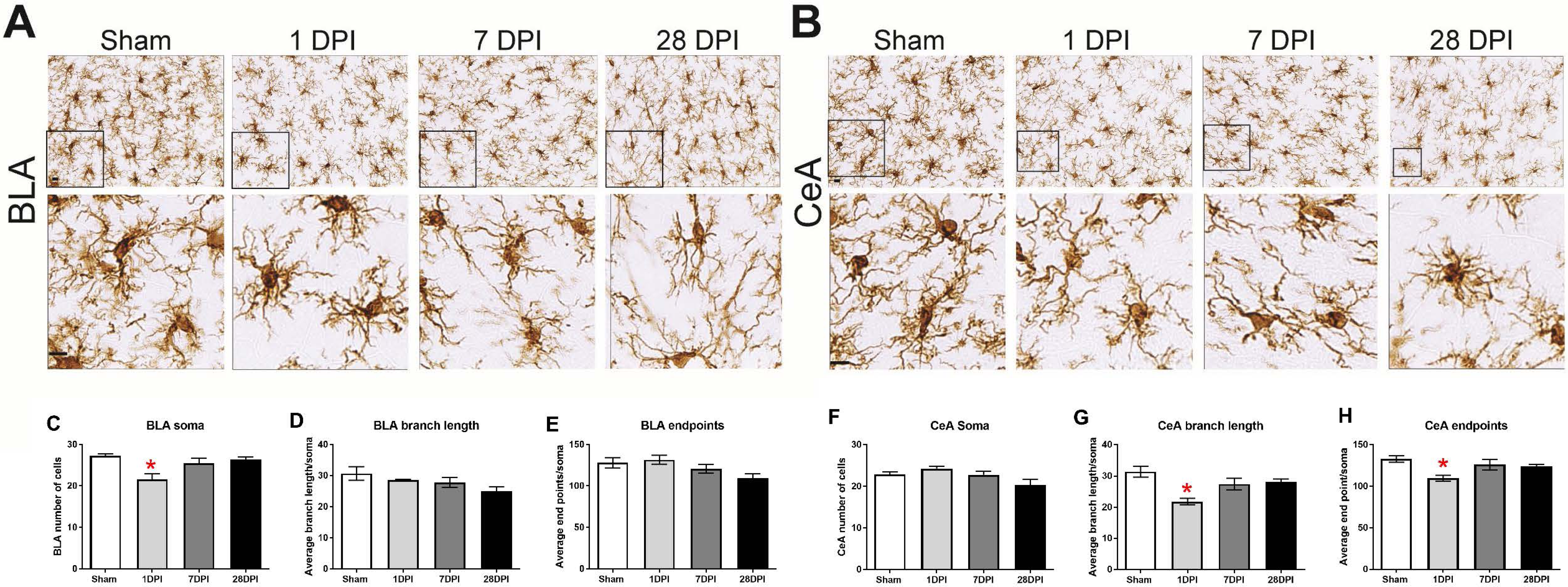
Acute, nuclei-specific microglial alterations occur following dTBI. Representative images of Iba1 staining in the **(A)** BLA and **(B)** CeA. Cropped photomicrographs correspond to the highlighted square. (Scale bars = 10µm). **C)** In the BLA, the number of microglia changed over time post-injury [One-way ANOVA F(3,8)=6.32; p<0.05; η^2^=0.70]. **D)** Branch length (μm/cell) or **E)** number of microglia process endpoints per cell did not significantly change over time [One-way ANOVA F(3,8)=2.32; p=0.15];[One-way ANOVA F(3,8)=3.18; p=0.09]. **F)** In the CeA, the average number of microglia per field did not significantly change over time [One-way ANOVA F(3,8)=3.17; p=0.09]. **G)** Contrasting to the BLA, branch length (μm/cell) [One-way ANOVA F(3,8)=5.27; p<0.05; η^2^=0.66] and **H)** number of microglia process endpoints per cell were significantly decreased at 1DPI in comparison to sham [One-way ANOVA F(3,8)=7.32; p<0.05; η^2^=0.73]. Bar graphs represents mean ± SEM. *n*=3 for each group

#### GFAP analysis

At 7 and 28 DPI, injured and sham rats were given a lethal dose of Euthasol^®^ and transcardially perfused with 4% PFA/PBS (sham=3, 7DPI=3, 28DPI=4). The brains were then cryoprotected in graded sucrose solutions (15%, 30%) and cryosectioned (20 µm). Sections were prepared exactly as previously described using rabbit anti-GFAP primary antibody (1:1000; Dako, catalog# Z0334) and biotinylated horse anti-rabbit secondary antibody (1:250; Vector Laboratories, Burlingame, CA, USA; Catalog#Ba-1100) then incubated with DAB ^20, 43^. The sections were dehydrated with ethanol and cleared in citrisolv before being cover-slipped with DPX.

Mosaic images encompassing both hemispheres were taken using a Zeiss microscope (Imager A2; Carl Zeiss, Jena, Germany) in bright-field mode with a digital camera. Quantitative analysis was performed using ImageJ Software (3.1.1v, NIH, Bethesda, MD, USA) on a Macintosh computer (OSX 10.11.6). The CeA was identified and traced based on topographical localization to the optic chiasm, rhinal fissure, and commissural stria terminalis. Grayscale digital images were then digitally thresholded to separate positive-stained pixels from unstained pixels. The percentage of argyrophilic (black) stained pixels was calculated for each image using the following formula:

Total area measured black/Total area measured × 100 =Percentage area with argyrophilic stain Five to six rat brain hemispheres were analyzed per rat (n=3-4/time point), depending on the quality of the section mounted (no folds or tearing within the area of interest). The same image was analyzed three times. The standard deviation between the percent area black was below 10% within each rat.

#### deOlmos silver stain analysis

Alternating sections of brains prepared by Neuroscience Associates Inc (Knoxville, TN) (see Iba1 staining methods) ^44^ were cryosectioned, mounted, stained using de Olmos aminocupric silver technique, counterstained with Neutral Red, and cover-slipped. Densitometric quantitative analysis was performed identical to GFAP analysis. Four hemispheres per rat (n=3/time point) were analyzed and each hemisphere was analyzed three times. These 12 statistical numbers indicating the percentage of black pixels were then averaged together to a single value (per rat) and subsequently used in statistical analysis. The standard deviation between the percent area black was below 10% within each rat.

#### Tissue dissection and protein extraction

At 28DPI, a new cohort of rats (n=5/each) were given a lethal dose of Euthasol^®^. Animals were transcardially perfused with ice-cold PBS for 3 minutes. The brain was rapidly removed and rinsed with ice-cold PBS. Tissue biopsies (1 mm diameter) taken bilaterally from the BLA and CeA were collected from 2 mm thick coronal sections made using a chilled rat brain matrix. Tissue biopsies were flash frozen and stored at −80°C until protein was extracted for automated capillary western analysis. Total protein was extracted from the BLA and CeA previously stored at −80°C. Tissues were homogenized in 250 μl of ice-cold extraction buffer (pH 8.0) containing 0.24 M Tris, 0.74 M NaCl, 100 μl TritonX100 with a protease inhibitor cocktail (complete, Roche Diagnostics; #11836153001). Tissue was homogenized with the Precellys^®^24 machine (Bertin Technologies, Montigny le Bretonneux, France) for 40 second bouts until the solution was completely clear (being chilled on iced for 2 minutes between bouts). Samples were then centrifuged at 3,000 × g for 15 minutes and the supernatant stored in 10-20 µl aliquots at −80C until analysis. Protein concentrations were determined using the Bicinchoninic acid assay (BCA) following manufacturer’s instructions (Pierce, Rockford, IL).

#### Automated capillary western –ProteinSimple (Wes)

Protein expression was evaluated using automated capillary western (ProteinSimple™, Biotechne, San Jose, CA). Experiments were run per manufacturer’s instructions and using products purchased through ProteinSimple™ (unless otherwise noted), including 12-230kDA capillary cartridges (SM-W004) and anti-rabbit detection modules (DM001). Proprietary mixtures included a biotinylated ladder and molecular weight standards as internal controls. Technical properties of the Simple Western^TM^ are described by Rustandi et al and Loughney et al ^45^. Prior to experiments, optimization was carried out for protein concentration, primary antibody concentration, multiplexing with a biological control (GAPDH), denaturing process, and exposure time (Table 1). Target protein concentration was chosen within the midpoint of the slope (0.125-2.5µg/capillary). Antibody concentration was based on evidence of antigen saturation at the chosen protein concentration. Target and GAPDH (loading control) were optimized for multiplexing to ensure chemiluminescence was within systems threshold and to ensure no protein-protein interactions. If chemiluminescent detection was above the threshold, secondary antibodies from Jackson ImmunoResearch Laboratories, Inc. (West Grove, PA) were used to dilute the manufacturer’s secondary antibody to reduce amplification of highly expressed targets (Goat Anti-Rabbit 111-005-045, HRP-Goat Anti-Rabbit 111-035-045, Goat Anti-Mouse 115-005-062, HRP-Goat Anti-Mouse 115-035-062). Denaturing temperature was chosen based on low signal to noise and baseline. The high-dynamic range of the exposures (algorithm in software) was used for data analysis in all experiments. Every capillary cartridge (25 capillaries) was run with the following controls: the same brain homogenate as a positive control, Erk as a system control, Antibody only, and protein only.

Protein extracts from amygdala samples were combined with sample buffer and master-mix (40 mM DTT, 0.1x ProteinSimple Sample Buffer, and 1x Fluorescent Standards) to achieve the desired protein concentration. Samples were then denatured via heating block at the optimized temperature (37°Cx30min). Duplicate capillaries of each sample were loaded per manufacturer’s instructions, the cartridge was centrifuged at 2500 RPM for 5 minutes, and placed into the automated capillary western machine where proteins were separated by size (electrophoresis), immobilized, and immunoprobed in individual capillaries. Once loaded into the instrument, the standard default metrics recommended by ProteinSimple were utilized for separation, incubation, and detection. The associated software, Compass (ProteinSimple^®^), generates an electropherogram with peaks corresponding to the expression of proteins of interest and calculates the area under the curve (AUC) for each peak (Figure 4E). To calculate relative protein expression, the AUC for the protein was normalized to the AUC for the biological control (GAPDH). The ratios from duplicate capillaries were averaged. Then, ratios from injured animals were normalized to shams in the same capillary cartridge.

**Figure 4:**
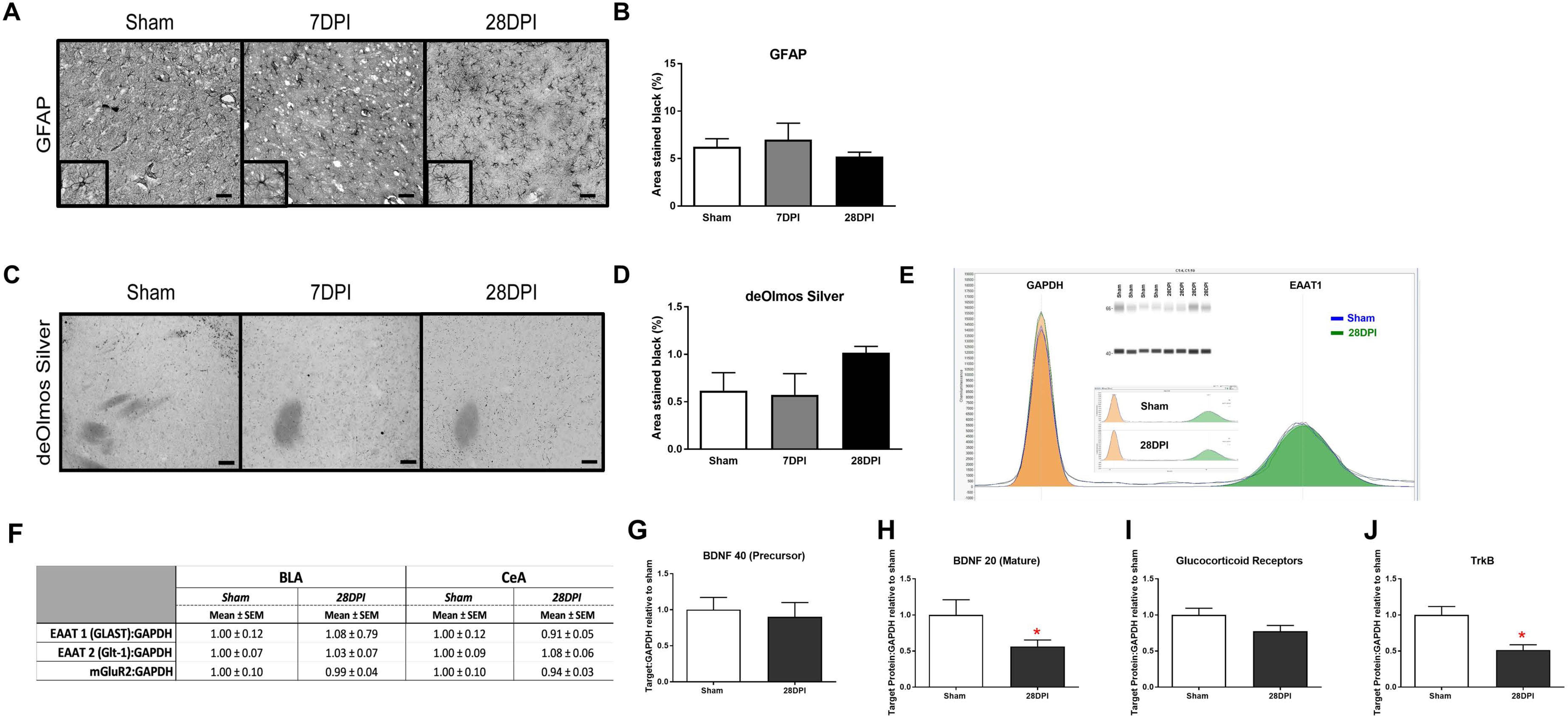
Absence of neuropathology, astrocytosis and molecular. **A)** Representative images of GFAP staining of the CeA (40× magnification; scale bar=50µm). **B)** Pixel density analysis of GFAP staining in the CeA revealed no significant differences following diffuse TBI [One-way ANOVA F(2,7)=0.80; p=0.49; *n*=3-4 per group]. **C)** de Olmos silver stained sections of the CeA (40× magnification; scale bar=50µm). **D)** Pixel density analysis of aminocupric silver stained tissue revealed that there was no difference following diffuse TBI. [One-way ANOVA F(2,6)=0.22; p=0.40; *n*=3-4 per group]. Automated capillary western blots (in duplicate) were performed to quantify the amount of synaptic glutamate transporters (Glt-1, GLAST) and pre-synaptic glutamate receptors (mGluR2, GluR) at 28DPI. **E)** Representative quantification of sham and injured GLAST (EAAT1) shown individually, overlapped and with recreated western blot. **F)** Levels of GLAST, Glt-1, and mGluR2 were similar to sham indicating an alternative mechanism for observed changes in glutamate neurotransmission (*n*=4-5 per group). **G)** The precursor protein to BDNF was similar to sham at 28DPI. **H)** Protein levels of mature BDNF were decreased by 43% (t_4,5_=2.07; p=0.04; *d*=1.33). **I)** GluR approached significance with a 20% decrease (t_4,5_=1.854; p=0.053). **J)** TrkB receptors were decreased by 49% (t_4,5_=3.68; p=0.004; *d*=2.41). *n*=4-5 per group. Bar graphs represent mean ± SEM.

#### Statistical analysis

The number of animals required to reach a 20% difference at 80% power at a significance level of 0.05 was completed based on previous studies by Thomas *et al* ^*32*^. All data were analyzed using a customized Microsoft Excel^®^ spreadsheet and Graphpad^®^ software. For values in which mean comparisons were used where the independent variable was nominal (Sham, 1DPI, 7DPI and 28DPI) and the dependent variable was continuous, a one-way ANOVA with Dunnet’s post-hoc comparison was conducted. For data involving dependent variables of ordinal nature, a Kruskal-Wallis with a Dunn’s post-hoc comparison was conducted. Parametrically analyzed data passed Shapiro-Wilks normality tests and Brown-Forsythe variability tests. Data not passing Shapiro-Wilks normality tests or Brown-Forsythe variability test were analyzed with a Kruskal-Wallis (KW) analysis and Dunn’s post-hoc comparison. Sham groups were the result of the combination of time points paralleling DPI as the groups were not statistically different from one another. Outliers for neurological assessment were determined using three standard deviations from the mean. Using these metrics, two rats were excluded from consideration (7-day sham, 7DPI). All data points for amperometry analysis were selected to represent each region of each rat using the dual, paired MEA recording sites and were averaged and used as a single data point. Outliers for amperometric analysis were determined using a ROUT test with Q = 5% (BLA=1 of 28; CeA=4 of 28). Additional analysis of extracellular clearance of glutamate in the CeA assessing for the potential effects of multiple additions of glutamate and differential effects of nuclei-specific microglial activation were conducted with a two-way ANOVA with a Tukey comparison. To be included in the amperometric recordings, the subject had to be included in the neurological assessment analysis. Molecular data for glutamate transporters were analyzed using an unpaired, two-tailed Student’s t-test. Based on our hypothesis of a decrease in protein levels for BDNF, TrkB, and GluR, we used a one-tailed Student’s t-test. For all instances, statistical significance was defined at p<0.05.

#### Effect size

For parametric calculation using the one-way ANOVA, eta squared η^2^ was conducted as described by Lakens 2013^46^. η^2^ was evaluated and reported such that 0.02 represents a small effect size, 0.13 represents a medium effect size and 0.16 represents a large effect size. Data comparing two means and analyzed using a Student’s t-test were evaluated using Cohen’s d. For nonparametric calculations using the Kruskal-Wallis test, Pearson’s r was reported as described by Fritz et al. 2012 ^47^. To do so, a Mann Whitney U post-hoc comparison was done between the two groups identified to have a difference as indicated by the Dunn’s post-hoc comparison. Then a Pearson’s r effect size calculation was evaluated and reported such that a 0.1 equals a small effect size, 0.3 represents a medium effect size and 0.5 represents a large effect size.

#### Data analysis validation

All outcome measures were evaluated for variability based on cohort membership, cage-mates, cage changes, time/order of open field testing, and time of recordings. Of these, no variables, other than injury status, could explain the variation observed in the presented data sets.

## Results

### dTBI induces the expression of anxiety-like behavior

Representative traces of first 5 minutes of open field exploration illustrate behavioral performance between brain-injured and sham rats (Figure 1B-C). Distances traveled at 7DPI and 28DPI were similar when compared to shams [F(2,29)=3.23; p=0.05; *n*=10-12; Figure 1D]. Further, rats at 7DPI did not significantly alter their duration in the center of the open field, whereas rats at 28DPI spent 57% less time in the center when compared to shams [KW=6.71; p<0.05; Figure 1E]. Rats at 28DPI also made 54% fewer entries into the center of the open field when compared to shams [KW=6.83, p<0.05; r=0.46; Figure 1F]. These data indicate that mFPI results in the expression of anxiety-like behavior by 28DPI.

### Glutamate neurotransmission in the BLA does not change over 1-month following dTBI

As the main sensory nuclei for anxiety-like behavior, the BLA was examined for alterations of glutamate neurotransmission ^48, 49^. BLA tissue was depolarized using a local application of isotonic 120 mM KCl (75-150 nL) to evoke the release of neurotransmitters, where the glutamate-sensitive MEA measured hydrogen peroxide as a surrogate for glutamate concentration. Similar volumes of KCl were applied to the extracellular space (Supplementary Figure 4A). KCl-evoked glutamate release in the BLA was not significantly different between 7DPI and 28DPI brain-injured rats when compared to shams [KW=4.63 p=0.10; Figure 2D].

Evaluation of BLA extracellular glutamate clearance parameters was accomplished through local application of 100 µM exogenous glutamate. There were no statistically significant differences in the volume of glutamate applied for each peak (Supplementary Figure 4B). Peaks were amplitude-matched to control for Michaelis-Menten kinetics, then analyzed for glutamate clearance parameters [F(2,21)=0.24; p=0.79]. Uptake rate was unaltered within the BLA at 7DPI or 28DPI following injury when compared to shams [F(2,21)=1.24; p=0.31; Figure 2E]. Furthermore, time taken for 80% of maximum applied glutamate to clear (T_80_) was not altered within the BLA at 7DPI or 28DPI when compared to shams [F(2,21)=0.140; p=0.870; Figure 2F]. These data indicate that evoked glutamate release and glutamate clearance parameters were not influenced by mFPI at 7DPI or 28DPI.

### Glutamate neurotransmission in the CeA is significantly altered over 1-month following dTBI

The CeA is the main amygdaloid nucleus for regulating anxiety-like behavior ^21, 48, 49^. Surrounding tissues, including pre-synaptic terminals along the BLA-CeA axis, were depolarized to evoke the release of glutamate using local application of 75-150 nL of 120 mM isotonic KCl. There were no statistically significant differences in the volume of applied KCl [Supplementary Figure 4C]. Representative traces of KCl-evoked glutamate release are shown in Figure 2G. Glutamate released in CeA was 50% less at 28DPI when compared to shams [F(2,19)=4.74; p<0.05; η^2^=0.33; Figure 2H].

Evaluation of CeA extracellular glutamate clearance was accomplished through local application of 100 µM exogenous glutamate. Representative traces of resulting glutamate peaks chosen for analysis are shown in Figure 2I. There were no statistically significant differences in the volume of glutamate applied for each peak [Supplementary Figure 4D]. Peaks were amplitude-matched to control for Michaelis-Menten kinetics, then analyzed for glutamate clearance parameters. Uptake rate was significantly decreased at 7DPI and 28DPI when compared to shams resulting in 40% and 58% decreases, respectively [F(2,22)=7.88; p<0.01; p<0.05 for 7DPI and p<0.01 for 28DPI; η^2^ = 0.42; Figure 2J]. Further, T_80_ of 28DPI rats was 43% longer when compared to shams [F(2,22)=5.00; p<0.05; η^2^ = 0.31; Figure 2K]. Uptake was independent of subsequent additions of local applications glutamate to the CeA [RM F(2,21)=8.16; p<0.01; Figure 2L]. These data indicate that evoked glutamate release and glutamate clearance parameters in the CeA were significantly influenced by mFPI over 28DPI.

### Nuclei-specific microglia deramification manifest acutely along BLA-CeA axis following dTBI

In the BLA, microglial cell count significantly decreased at 1DPI [F(3,8)=6.32; p<0.05; Figure 3C]. The branch length per cell (μm/cell) did not significantly change over time [F(3,8)=2.32; p=0.15; η^2^=0.70; Figure 3D]. Furthermore, the number of microglia process endpoints per cell did not significantly change over time [F(3,8)=3.18; p=0.09; Figure 3E].

Identical analysis was performed on microglia in the CeA. The microglial cell count per field did not significantly change over time [F(3,8)=3.17; p=0.09; Figure 3F]. The branch length per cell (μm/cell) significantly decreased at 1DPI [F(3,8)=7.32; p<0.05; η^2^=0.73; Figure 3G]. The number of microglia process endpoints per cell was also significantly decreased at 1DPI [F(3,8)=5.27; p<0.05; η^2^=0.66; Figure 3H].

The same three variables were evaluated with a two-way ANOVA to determine that the influence of time-post injury was similar in the BLA and CeA. All 3 outcome measures indicated an interaction that differed depending on the nucleus being evaluated. The number of cells (F(3,16)=7.85; p<0.05), endpoints (F(3,16)=5.152; p<0.05), and process length (F(3,16)=3.94; p<0.05] all differed between regions, indicating that TBI-induced changes in microglial morphology at 1 and 28DPI are region dependent (Supplementary Figure 5).

**Figure 5:**
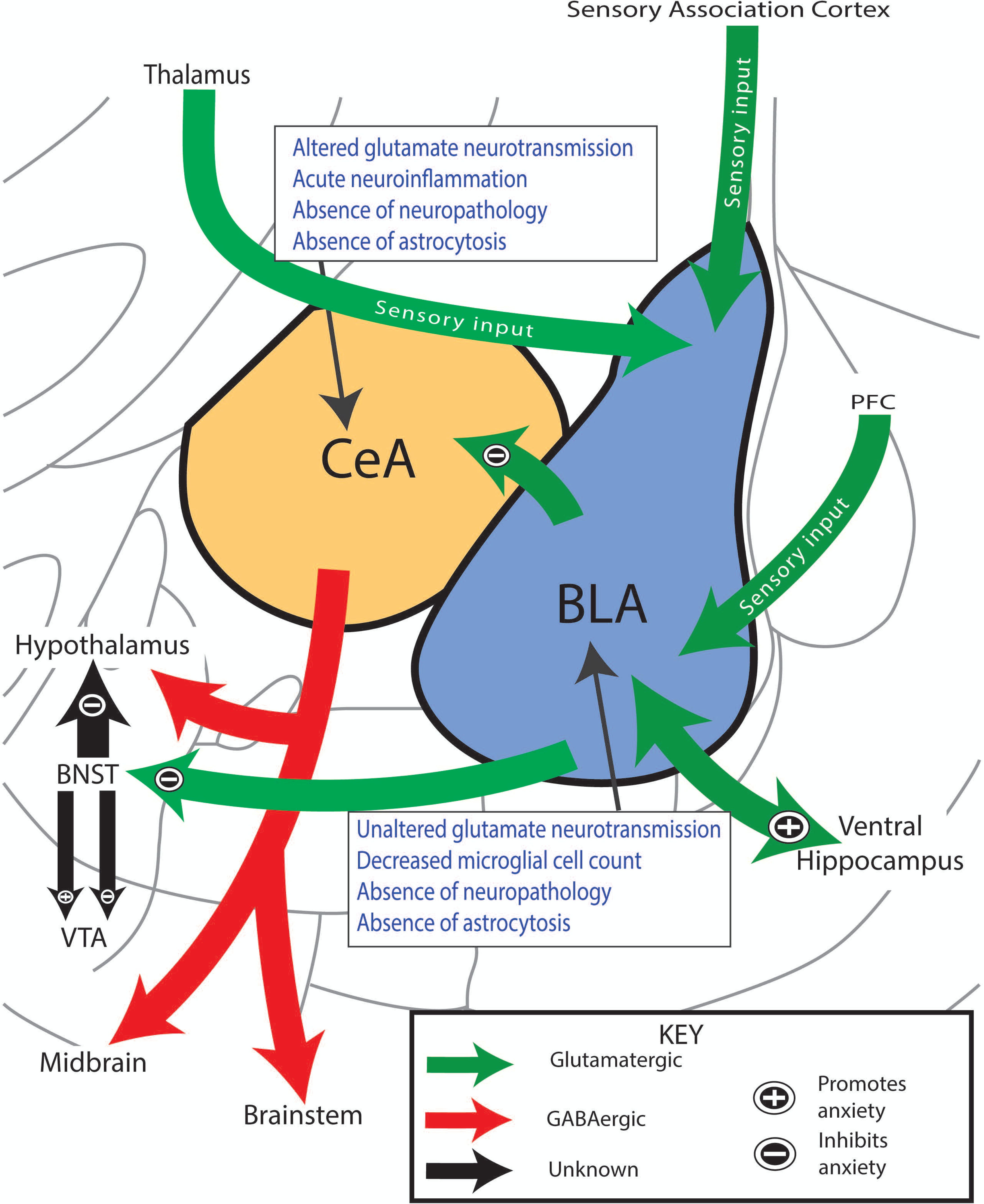
Anxiety-like behavior coincides with altered glutamate neurotransmission in amygdala circuitry. Simplified circuitry of the rat’s efferent and afferent projections of the amygdala that are considered to play important roles in the regulation of anxiety-like behavior. Glutamatergic projections enter the BLA from the PFC and thalamus and carry sensory information of the overall anxious state of the rat. The BLA interprets this information and sends efferent outputs to either promote (+) or inhibit (-) the display of anxiety-like behavior. Activation of BLA-CeA circuitry has shown to mediate anxiolytic behavior. Boxes summarize findings. PFC: prefrontal cortex; BLA: basolateral amygdala; CeA: central nucleus of the amygdala; BNST: bed nucleus stria terminalis; VTA: ventral tegmental area

### Absence of activated astrocytes over 1 month after dTBI

Robust astrocytosis in the BLA has been previously reported to be present at 7DPI and to resolve by 28DPI following experimental dTBI using mFPI ^20^. Astrocytes may contribute to glutamate neurotransmission due to their role in clearance of glutamate from the extracellular space, and thus were analyzed using GFAP staining in the CeA. There was no evidence that mFPI influenced astrocytosis over time in the CeA [F(2,7)=0.80; p=0.49; Figure 4A-B].

### Absence of overt neuropathology in the CeA following dTBI

The BLA has been described as lacking overt neuropathology at 7DPI and 28DPI ^20^, however, the CeA has not been evaluated in a model of mFPI. Using de Olmos silver stain technique, neuropathology in the CeA was assessed at 7DPI and 28DPI compared to shams. No overt pathology was found to occur in the CeA following mFPI [F(2,6)=1.99; p=0.22; Figure 4C-D].

### Alteration in glutamate neurotransmission independent of glutamate transporters and pre-synaptic receptors

Observed alterations to glutamate neurotransmission could result from altered levels of glutamate transporters or presynaptic metabotropic receptors ^50^. Proteins involved in glutamate transport (EAAT1(GLAST) EAAT2 (Glt-1)) were evaluated at 28DPI for the BLA and CeA. Levels of GLAST and Glt-1 were similar to sham [GLAST (BLA t_8_=0.59 p=0.57; CeA n=4, t_7_=0.75; p=0.49; Glt-1 (BLA t_7_=0.25; p=0.81; CeA t_6_=0.81; p=0.45)];] (Figure 4E-F). Pre-synaptic metabotropic glutamatergic receptors (mGluR2) levels did not change in comparison to shams (BLA t_9_=0.09; p=0.93; CeA t_6_=0.51; p=0.63;) (Figure 4F), indicating that observed changes in glutamate neurotransmission were independent of changes in total protein levels of glutamate transporters and mGluR2. Supplementary Figure 6 depicts localization of these molecules on a presynaptic terminal.

### Neuropsychiatric molecular targets decreased at 1-month post-dTBI

As BDNF, GluR, and TrkB, have been shown to influence presynaptic glutamate release, we tested the hypothesis that dTBI caused decreased levels of BDNF, GluR, and TrkBin the amygdala by 28DPI ^51^. Protein levels of mature BDNF were decreased by 43% (t_7_=2.07; p=0.038; *d*=1.33; Figure 4H) (BDNF precursor was not different) and GluR approached significance with a 20% decrease (t_7_=1.854; p=0.053; Figure 4I). TrkB receptors were decreased by 49% (t_7_=3.68; p=0.004; *d*=2.41; Figure 4J). These data indicate that the neuropsychiatric molecular targets, BDNF and TrkB, were decreased in brain-injured rats at 1-month post-injury coinciding with the expression of anxiety-like behavior and changes in evoked glutamate neurotransmission.

## Discussion

Multifactorial mental illnesses are the result of poorly understood pathophysiology that confounds treatment approaches leading to poor symptom control. These are the first experiments that demonstrate dTBI initiates a cascade of molecular events in the amygdala capable of contributing to the expression of anxiety-like behavior. By one-month post-injury, decreased glutamate release and slower glutamate clearance within the CeA coincided with the expression of anxiety-like behavior. There were no changes in the BLA. Changes in glutamate neurotransmission occurred despite similar protein levels of glutamate transporters, mGluR2, and in the absence of overt neuropathology or astrocytosis. BDNF and TrkB protein levels were significantly decreased by 28DPI, indicating a potential role for BDNF/TrkB signaling in decreased glutamate neurotransmission. Early region-dependent neuroinflammation precedes changes in neurotransmission. These data indicate novel targets in the quest for understanding the pathophysiology of TBI-induced anxiety.

Characterized by validated metrics reviewed by Gould *et al.* ^33^, we show late-onset anxiety-like behavior following dTBI. By 28DPI, rats spent less time and made fewer entries into the center of the open field when compared to shams, indicative of anxiety-like behavior. Distance traveled between shams and brain-injured rats were similar and therefore not a confounding variable for anxiety-like behavior. Limbic system circuitry has been shown to regulate affective symptomatology during neurological assessment tasks such as open field ^21, 22, 52^. The glutamatergic connections between the BLA and CeA have a robust influence in the expression of affective behavior in rodents, integrating input from numerous regions of the brain (Figure 5) ^21, 22, 52, 53^. In short, sensory information transfers to the BLA from the prefrontal cortex (PFC), thalamus, and sensory association cortex. From here, the BLA sends projections to the ventral hippocampus, bed nucleus of the stria terminalis (BNST), or CeA that either promote or extinguish anxiety-like behavior ^22, 48, 53, 54^. The BLA-CeA circuitry is known to be involved in the production of affective behaviors ^48, 49^; however, the functional, structural, and molecular components underlying the development after TBI have never been evaluated.

For real-time recordings of extracellular neurotransmission, *in vivo* amperometry coupled with glutamate selective multielectrode arrays were utilized for their excellent spatial and temporal resolution and the presence of sentinel channels to verify glutamate specificity. Localized KCl administration induced non-specific neurotransmitter release in which glutamate-sensitive MEA’s measured synaptic glutamate overflow. In the BLA, no significant changes were detected, however, in the CeA glutamate release was decreased by 28DPI compared to uninjured shams. Reduced glutamate release indicates less stores available for potential neuronal communication between those located in presynaptic neurons, interneurons, and astrocytic processes. The second peak within the biphasic profile of KCl-evoked glutamate release (Figure 2G) provides evidence that glial transmission could be contributing to glutamate overflow whereas the first peak is thought to primarily be the response of presynaptic neurons ^55, 56^. While the amplitude of the first peaks was significantly decreased at 28DPI compared to shams (data not shown), area under the curve was used to evaluate all aspects of glutamate neurotransmission. Changes in glutamate overflow could arise from impaired vesicular loading ^57^ or secondary to inhibitory GABA-ergic modulation ^58^. We did not find an injury-induced change in the protein level of mGluR2 in the CeA, indicating that total mGluR2 is not predictive of changes in presynaptic glutamate release. Decreased BDNF/TrkB/GluR protein levels can reduce presynaptic glutamate release, however, localization, cell type, and functional studies are necessary to identify their role ^29, 30^. We also did not observe astrocytosis or perturbation of microglia at 28DPI, further indicating a role for the BDNF/TrkB/GluR signaling cascade. Decreased glutamate release in the CeA may be indicative of inhibition of the BLA-CeA circuit, which coincides with increased anxiety-like behavior ^21^. Thus, while novel in TBI, our results parallel the consensus that dysregulation of the CeA may be concomitant with the expression of affective conditions ^59, 60^.

Locally applied glutamate to the BLA and CeA allowed us to calculate glutamate clearance from the extracellular space, primarily mediated by glutamate transporters (Glt-1 and GLAST) located on adjacent astrocytes in the rodent ^27^. The affinity of glutamate transporters can be evaluated using glutamate uptake rate, whereas the number of membrane bound transporters is estimated using T_80_ ^61^. In the BLA, glutamate clearance parameters did not change as a function of injury over time. In the CeA, the uptake rate was significantly slower at 7DPI and 28DPI while glutamate took significantly longer to clear (T_80_) by 28DPI. For calculations of glutamate parameters, three consecutive peaks in each region were analyzed. The consecutive peaks were reproducible, and the uptake rate remained significantly different regardless of the repeated applications, confirming that observed alterations to glutamate clearance are not caused by immediate tissue compensation. Changes in glutamate clearance were also not due to changes in the protein levels of glutamate transporters. Slower glutamate clearance could also be caused by surface expression, post-translational modifications, or adaption of glia and adjacent cells ^62, 63^.

No significant alterations were detected in the BLA in anesthetized rats in this study, although studies using focal TBI models indicate that BLA circuitry becomes weakened through altered neuronal excitability, changes in N-methyl-D-aspartate (NMDA) receptors, and changes to GABAergic production proteins (GAD-67), all of which may provide compensatory responses that primarily influence glutamate release in the CeA ^20, 64, 65^. Previously, we reported increased neuropathology in the somatosensory cortex and thalamus following dTBI, which could also alter sensory input into the BLA ^66, 67^. Assessment of glutamate neurotransmission in both the BLA and CeA of awake-behaving rats is needed to evaluate simultaneous glutamate signaling in direct response to anxiety-inducing tasks. This model would provide the ability to test the effectiveness of anxiolytic intervention in mediating glutamate signaling within the CeA as a universal consequence of dTBI.

Morphological analysis of microglia in the BLA and CeA nuclei revealed a nuclei-dependent neuroinflammatory response that preceded changes in glutamatergic neurotransmission along the BLA-CeA circuit. Deramified microglia were identified in the CeA, but not the BLA, at 1DPI. In the BLA, microglial cell counts decreased by 1DPI without evidence of microglial de-ramification, suggesting that dTBI does not lead to a neuroinflammatory response despite evidence of perturbation. In the CeA, microglial cell counts did not change over time, while the summed process length and number of microglial process endpoints per cell were found to be significantly decreased at 1DPI. These data suggest that microglia become de-ramified in the CeA early and recover to sham levels by 7DPI, indicative of an acute inflammatory response after injury ^68^. Subsequently, we analyzed CeA neuropathology to determine if damaged neurons instigated the microglial response in the CeA. However, in accordance with previous results published for the BLA ^20^, silver staining did not reveal any evidence of overt neuropathology. Analysis of astrocytosis revealed that in contrast to our previous work in the BLA, astrocytes were not activated in the CeA following dTBI ^20^. This indicates that CeA microglial activation is independent of the presence of pathology and may play an alternative role in the sequalae of events after TBI ^69^. Chronologically, dTBI initiates an immediate release of glutamate, a stress response that includes increased circulating corticosterone levels, and a cascade of inflammatory cytokines that could mediate CeA microglial activation through ionotropic and metabotropic glutamate receptors and glucocorticoid receptors ^25, 70^. In comparison to the development of affective disorders by other etiologies, psychosocial stress-induced neuroinflammatory responses have been proposed as the etiology, with repeated social defeat being indicated to alter microglia morphology in the amygdala ^71^. Together, this indicates that TBI-induced microglial activation is mediating circuit reorganization in the amygdala similar to that observed in the development of affective disorders of alternative idiopathic etiologies.

We have previously reported a significant decrease in basal plasma corticosterone levels and a blunted stress response at 56DPI, indicating chronic dysregulation of the HPA-axis in this TBI model ^43^. Chronic dysregulation of corticosterone in response to stress has also been supported in other TBI models at both earlier and later time points^72, 73^. The central amygdala mediates the HPA-axis response to stress, where changes in circulating corticosterone levels, GR, BDNF, and Trk-B receptor expression have been implicated in the pathology of affective disorders (anxiety-like behavior and posttraumatic stress disorder-like phenotypes) in addition to presynaptic glutamate signaling ^30, 51, 74^. Further evaluation is necessary to determine whether a role exists for corticosterone dysregulation and corticosterone regulated receptors in the development of anxiety-like behavior and changes in glutamate neurotransmission over time following dTBI.

Clinically, it is difficult to differentiate the initiating insult for patients experiencing affective symptoms, such as anxiety or PTSD, as similar pathophysiology exists between dTBI and chronic stress ^75-78^. This study provides a novel link between late-onset anxiety-like behavior and altered glutamate neurotransmission within the CeA, that parallels clinical studies that implicate dysfunctional amygdala processing in the expression of affective symptomatology post-injury ^14, 15, 54, 79-81^. Decreased BDNF/TrkB protein levels implicate one pathway by which dTBI can influence anxiety-like symptoms. Identification of common pathways for the development of TBI-induced and non-TBI-induced affective disorders are instrumental in treatment of symptoms, rehabilitation guidelines, and implementation of clinical approaches.

## Acknowledgements

The authors would like to thank the following individuals for their assistance on these projects. Dr. Kimbal Cooper at Midwestern University for statistical guidance. Ms. Katherine R. Giordano for technical assistance and microscopy optimization. Ms. Carol Haussler, Samantha W. Ridgway, and Dr. Gokul Krishna for valuable editing and feedback. Research reported in this publication was supported in part by National Institute of Neurological Disorders and Stroke of the National Institutes of Health under Award Number R01NS100793 awarded to TCT, Valley Research Partnership (VRP) P1 (P1201607) awarded to JAB, Phoenix Children’s Hospital (PCH) Leadership Circle Grant awarded to TCT, PCH Mission Support Funds awarded to TCT, Arizona Biomedical Research Commission through Arizona Department of Health Services (ADHS14-00003606) awarded to TCT, Director’s Research and Education fund at PCH Foundation awarded to TCT, Midwestern University ORSP and Midwestern Biomedical Sciences Department funds awarded to JAB. The content is solely the responsibility of the authors and does not necessarily represent the official views of the National Institutes of Health.

## Supplementary Figures Legends

**Supplementary Figure 1: Schematic of glutamate sensitive MEA**

Ascorbic acid and other aromatic compounds are repelled from the MEA due to the electroplated 1,3-phenylenediamine (mPD) exclusion layer. Glutamate sensing (bottom pair) and sentinel (top pair) recording sites are coated with BSA and glutaraldehyde. Glutamate oxidase is added to signal sites only to convert glutamate to α-ketoglutarate and peroxide. Peroxide is oxidized on the platinum recording sites as a result of an applied potential of 0.7 V vs. Ag/AgCl reference electrode, thereby producing a current that is correlated with the concentration of glutamate present. Sentinel sites (top pair) are unable to record glutamate due to the lack of glutamate oxidase. Top left corner depicts an MEA. Schematic and MEA are not to scale.

**Supplementary Figure 2: Representative confirmation of MEA placement**

Following amperometric experiments, brains were harvested and fixed for MEA placement. Brains were cut coronally at 40 µm. Here, a representative image of MEA placement into the anatomical ventral lateral hemisphere of the rat is shown. The black arrows indicate the outer track (BLA) and the inner track (CeA).

**Supplementary Figure 3: Skeletal Analysis Technique**

Individual steps for skeleton analysis of microglia morphologies of Iba1 stained tissue. Original photomicrographs were subjected to a series of uniform ImageJ plugin protocols prior to conversion to binary images which were then skeletonized. An overlay of a resulting skeletonized image (in green) and original photomicrograph shows the relationship between skeleton and photomicrograph. All skeleton analysis was completed on full sized photomicrographs (40× magnification; scale bar = 10µm).

**Supplementary Figure 4: Controls for applied solutions in amperometric recordings**

Anesthetized *in vivo* amperometric recordings of glutamate neurotransmission were conducted in sham, 7DPI, and 28DPI rats in the BLA (A/P: −2.4 M/L ± 5.1 D/V -8.0) or CeA (A/P: −2.4 M/L ± 3.9 D/V −8.0). Local applications of 120 mM potassium chloride solution (KCl) were made to depolarize surrounding neurons. No differences in the volume locally applied was observed between sham and brain-injured rats in the **A)** BLA [One-way ANOVA F(2,25)=0.47; p=0.63; *n*=8-12] or **C)** CeA [One-way ANOVA F(2,18)=0.74; p=0.49; *n*=6-9]. Local applications of exogenous 100 µM were made to measure extracellular glutamate clearance. All applied peaks were amplitude matched prior to analysis in the **B)** BLA [One-way ANOVA F(2,21)=0.24; p=0.79; *n*=6-10] and **D)** CeA[One-way ANOVA F(2,22)=0.05; p=0.37; *n*=7-10] to control for Michaelis Menten kinetics. Bar graphs represent the mean ± SEM.

**Supplementary Figure 5: BLA and CeA microglia respond differently to dTBI over time**

Microglial morphology changes as a function of time post-injury and region. (**A)** Microglial cell counts were different than sham after 1, 7 and 28DPI in the BLA [Two-way ANOVA F(3,16)=7.85; p<0.05] (**B**). The number of microglial process endpoints/cell was lower than sham after mFPI in the CeA [(3,16)=5.152; p<0.05]. (**C**). The summed process length/cell were lower than sham after mFPI in the CeA [F(3,16)=3.94; p<0.05]. Bar graphs represent mean ± SEM, *n*=3/group.

**Supplementary Figure 6: Representation of glutamatergic targets for protein quantification studies.** Representation of a glutamatergic synapse and the proteins that regulate glutamate neurotransmission. Diagram modified from Thomas et al. 2012 ^32^.

