## Supplementary Figures for "Diffuse TBI-induced expression of anxiety-like behavior coincides with altered glutamatergic function, TrkB protein levels and region-dependent pathophysiology in amygdala circuitry"

### Supplementary Figure 1

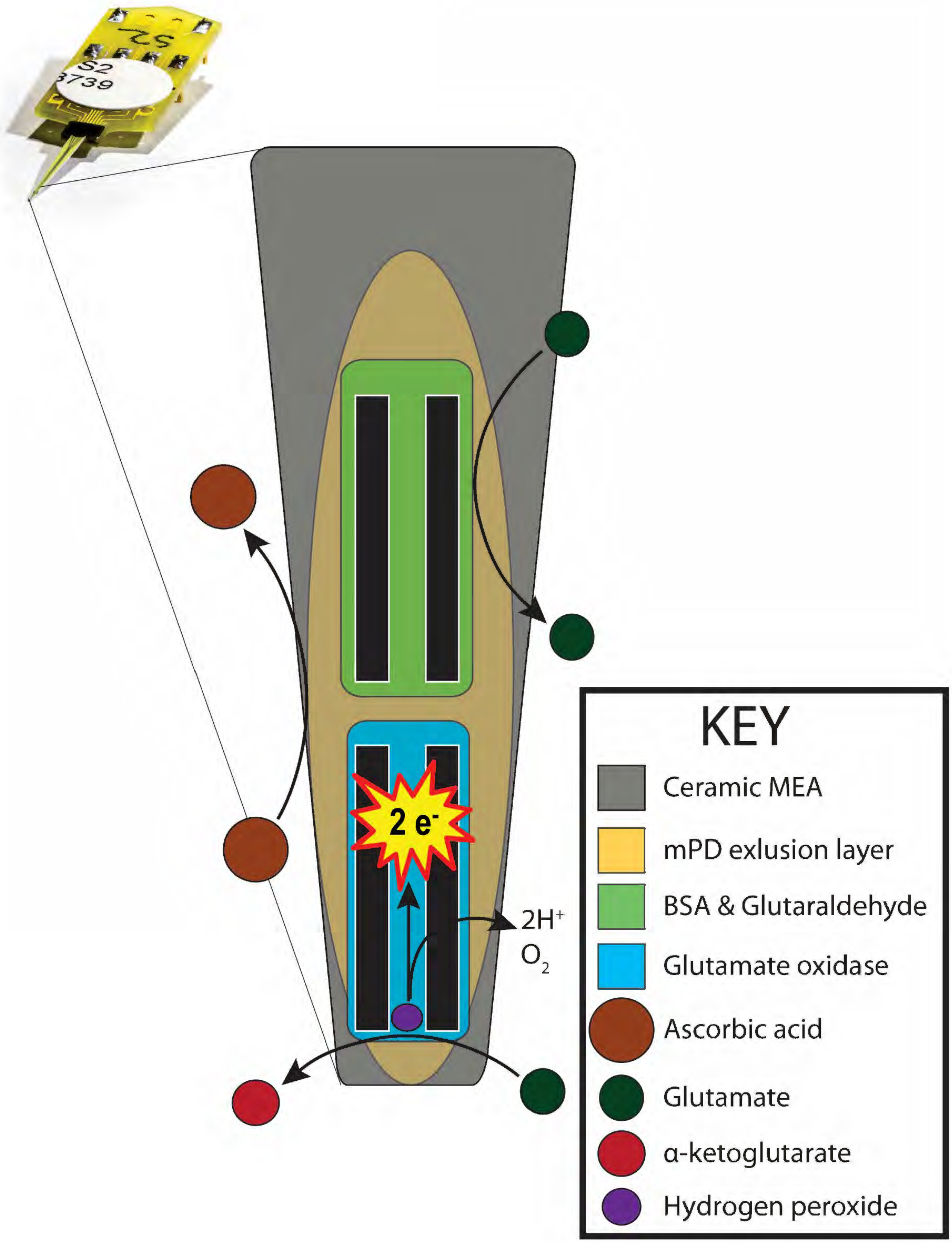

### Supplementary Figure 2

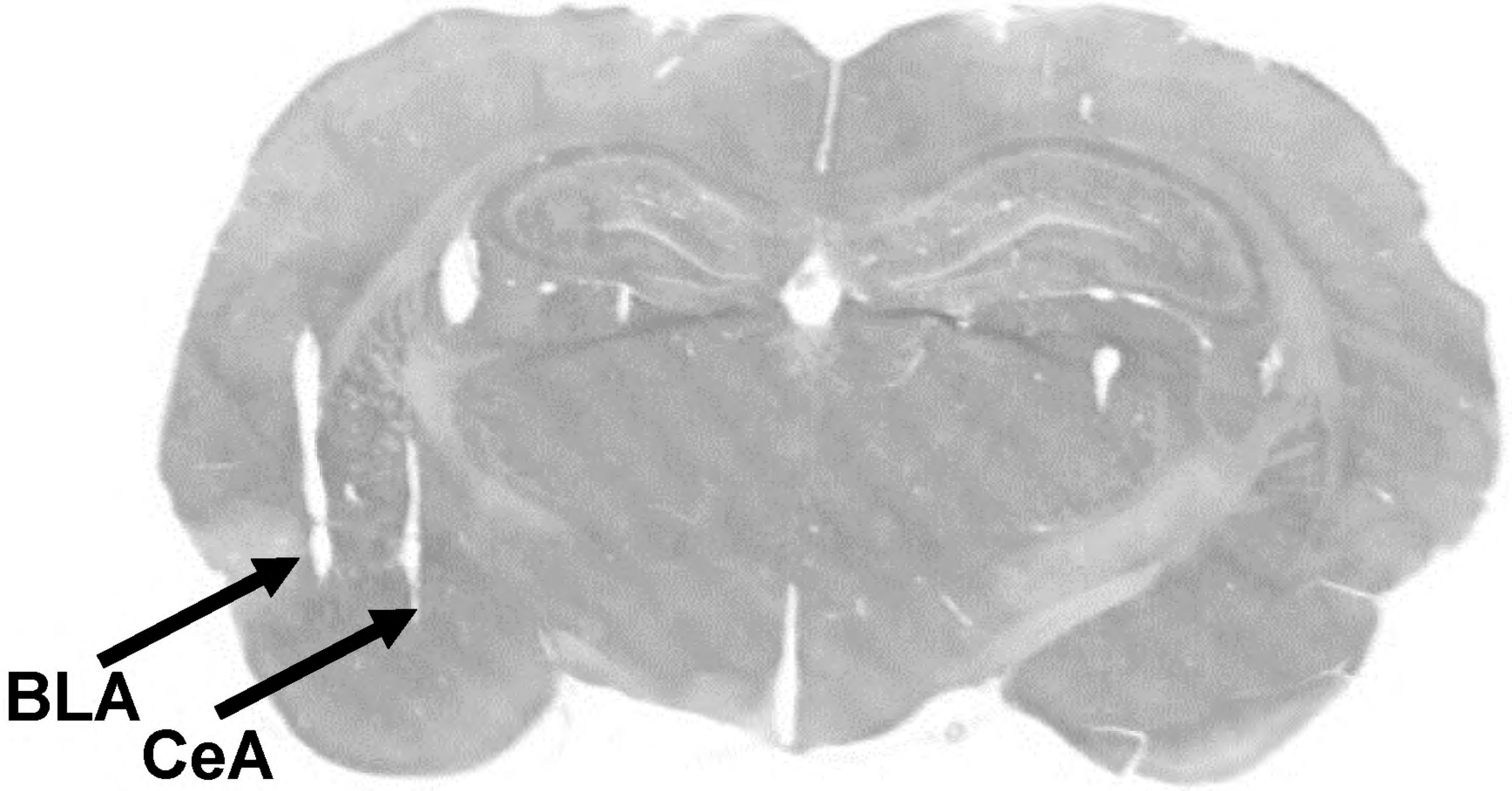

**Supplementary Figure 3**

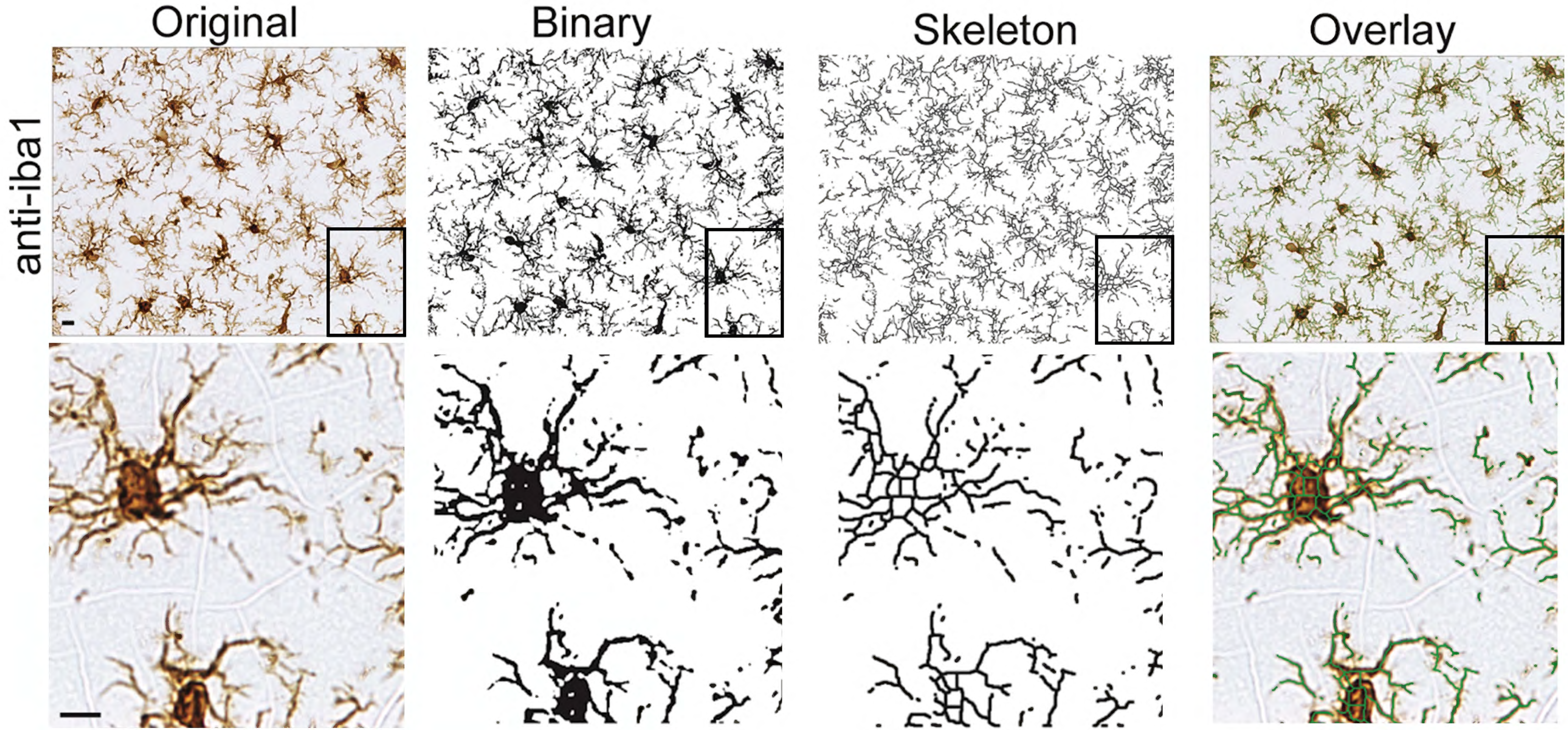

### Supplementary Figure 4

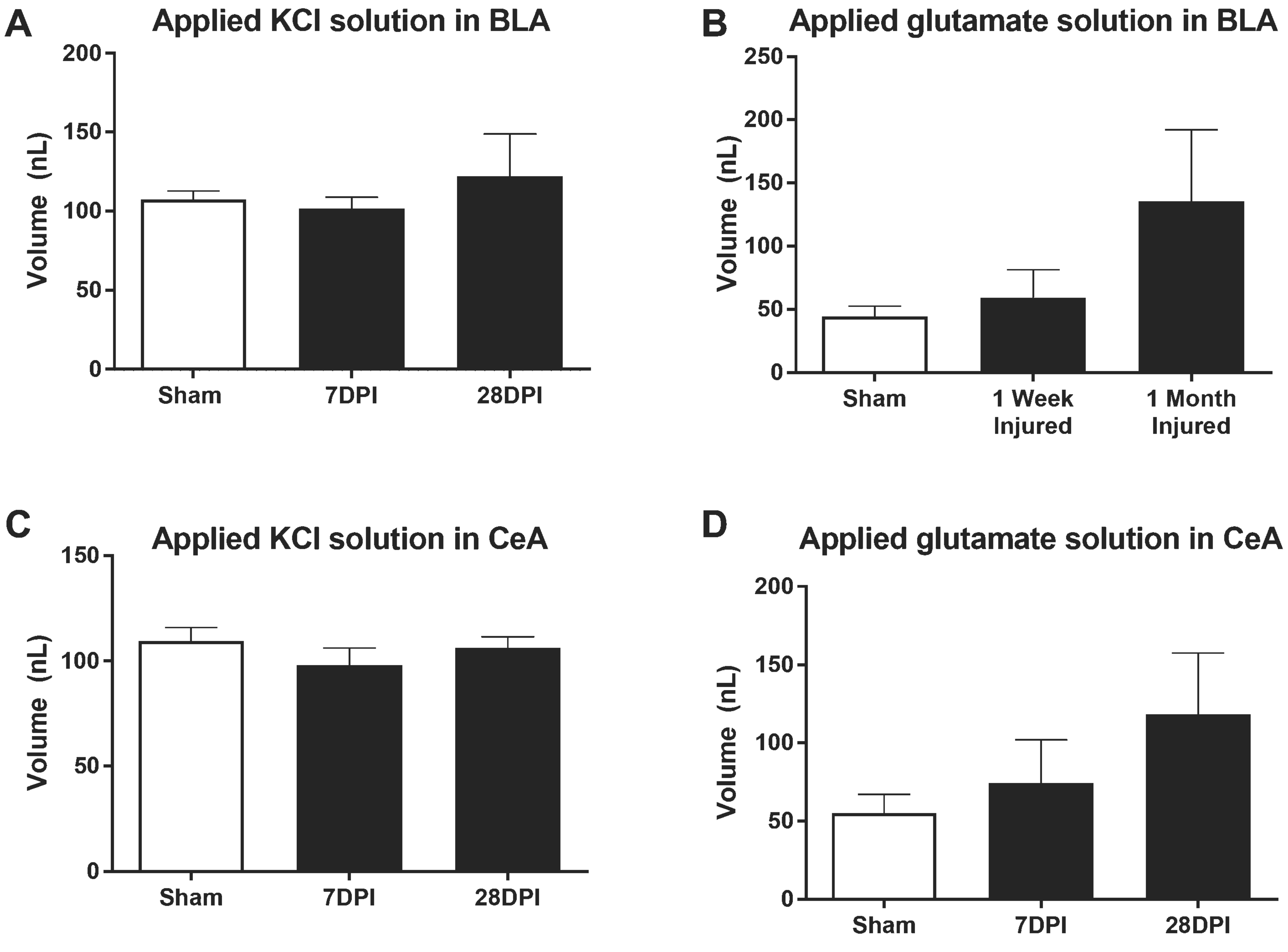

### Supplementary Figure 5

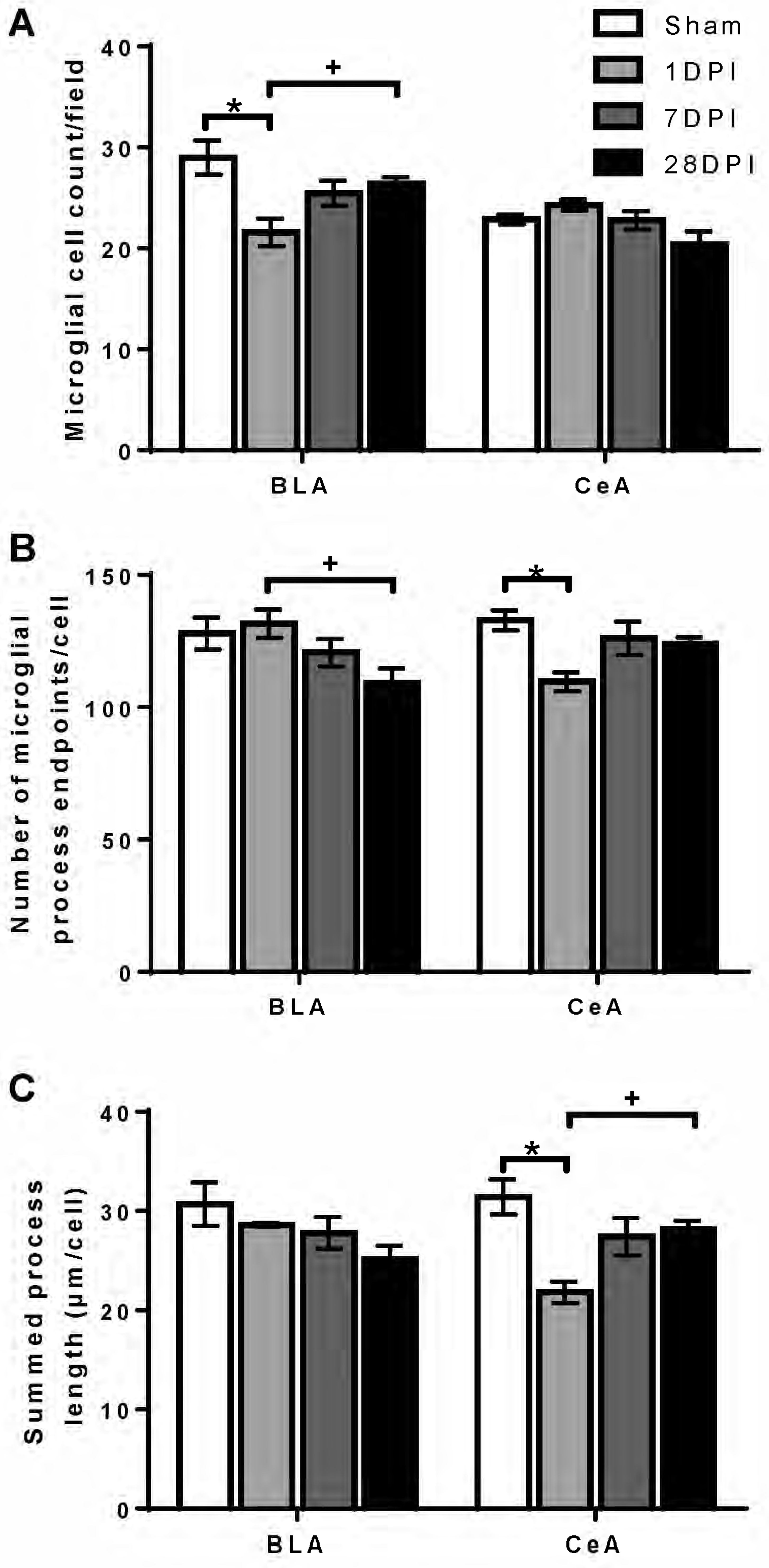

### Supplementary Figure 6

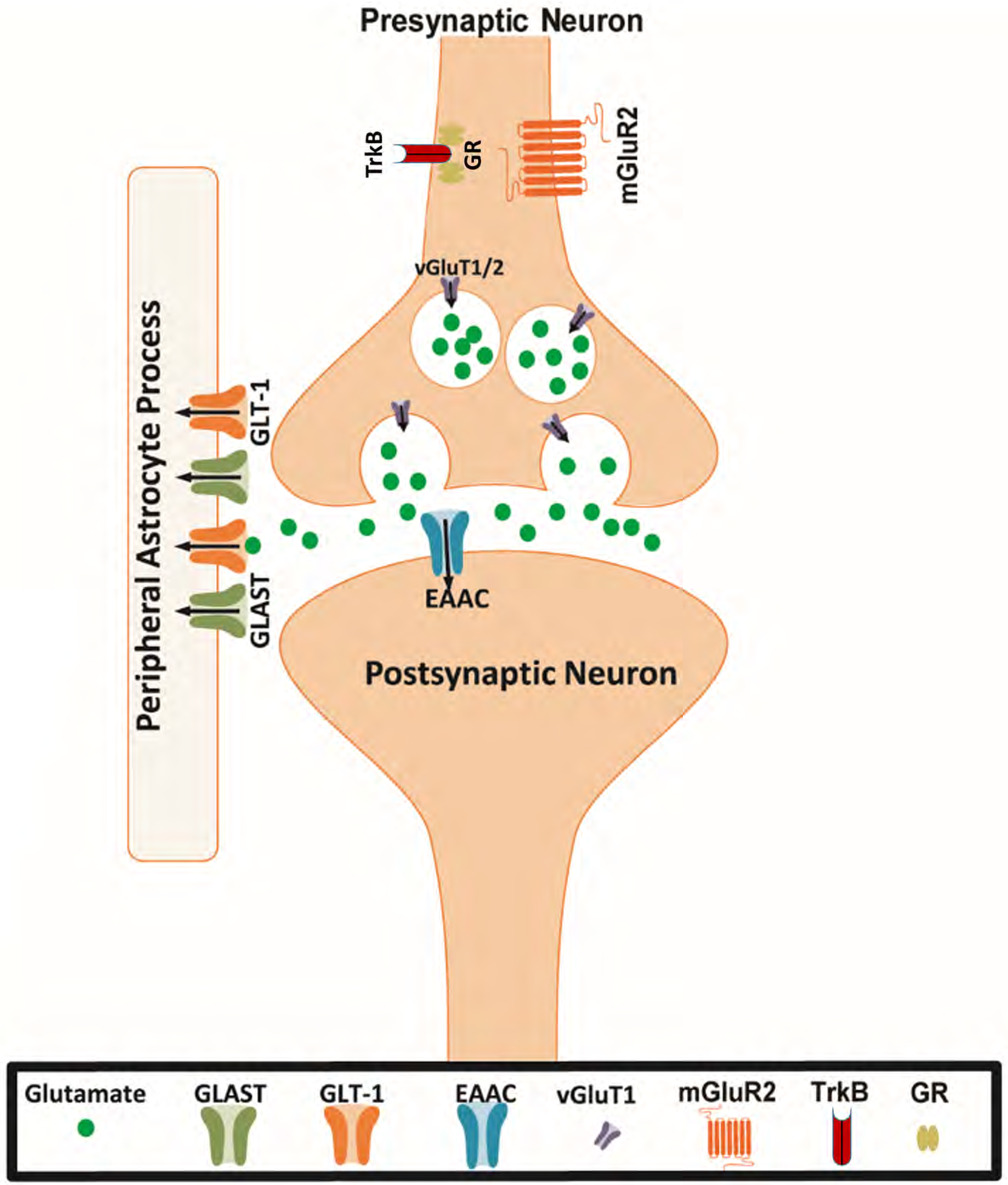

| Antibody | Species | Company, catalog number | Denaturing Temp (celcius) | [Protein] μg/μl | [Ig] | Biological control |
|---|---|---|---|---|---|---|
| BDNF | Ms | Abcam, ab108319 | 37Cx30min | 0.5μg/μl | 1:50 | GAPDH |
| GLUR/NR3C1 | Rb | Neo, A2164 | 37Cx30min | 0.5μg/μl | 1:25 | GAPDH |
| TrkB | Rb | Abcam, ab18987 | 37Cx30min | 0.5μg/μl | 1:100 | GAPDH |
| GLT-1 (EAAT2) | Rb | Abcam, ab205248 | 37Cx30min | 0.1μg/μl | 1:25 | GAPDH |
| Glast (EAAT1) | Rb | Abcam, ab181036 | 37Cx30min | 0.5μg/μl | 1:25 | GAPDH |
| mGluR2 | Rb | Abcam, ab150387 | 37Cx30min | 0.1μg/μl | 1:100 | GAPDH |
| GAPDH | Ms | Abcam, ab8245 | 37Cx30min | 0.5μg/μl | 1:25 | |
